# β-catenin Dynamics in Regenerating *Hydra*

**DOI:** 10.64898/2026.09.21.752673

**Authors:** Liora Garion, Yael Ascoli-Abbina, Noam Dori, Iris Pasvinter, Kinneret Keren

**Affiliations:** Department of Physics, Technion – Israel Institute of Technology, Haifa 3200003, Israel; Network Biology Research Laboratories and Russell Berrie Nanotechnology Institute, Technion – Israel Institute of Technology, Haifa 3200003, Israel

## Abstract

Body axis specification is essential for morphogenesis, yet how axial patterning is dynamically coordinated remains unclear. In *Hydra*, Wnt/β-catenin signaling is central to axial patterning, but its spatiotemporal dynamics are not well understood. Since nuclear β-catenin mediates Wnt-dependent transcription, its localization provides a relevant readout for pathway dynamics. Here, we track β-catenin abundance and nuclear localization *in vivo* during regeneration from bisected *Hydra*, tissue rings, and small tissue fragments. Following bisection, nuclear β-catenin is detected exclusively at the regenerating oral end, where a broad signal appears and then declines before focusing at the future head. Tissue rings, lacking both head and foot, exhibit similar oral-aboral asymmetry. In regenerating fragments, the initial response often spans nearly the entire tissue, including the future foot region, followed by an additional peak before stabilizing at the future head. Overall, our results show that the spatial restriction of nuclear β-catenin correlates with the divergence of regenerative dynamics toward head or foot formation, while the trajectories leading to these outcomes vary across different initial tissue configurations.

## Introduction

Establishment of the body axes is a fundamental step in organizing the body plan during animal morphogenesis. Across the animal kingdom, a small number of conserved signaling pathways are consistently implicated in axial patterning (Barresi and Gilbert 2024). The Wnt/β-catenin pathway is a prominent example governing key aspects of primary body axis formation in a range of animals, including mammals, amphibians, planarians and cnidarians, and its perturbation leads to abnormalities in axis specification (C. P. Petersen and Reddien 2009; Rim et al. 2022).

β-catenin is the central intracellular effector in the Wnt/β-catenin pathway, also known as the canonical Wnt pathway, linking extracellular Wnt signals to transcriptional responses (Rim et al. 2022). In the absence of Wnt signaling, a GSK3-containing destruction complex targets cytoplasmic β-catenin for degradation. Wnt signaling inhibits this process, allowing β-catenin to accumulate and enter the nucleus, where it associates with the TCF transcription factor to activate transcription of Wnt target genes. Importantly, β-catenin is also an essential structural component of cell–cell junctions. Since β-catenin functions both in cell–cell adhesion and in signaling, its partitioning among junctional, cytoplasmic, and nuclear pools creates the possibility of competition or coupling between these functions (Fagotto 2013; van der Wal and van Amerongen 2020). Consequently, changes in β-catenin transcription or overall protein abundance do not necessarily reflect corresponding changes in canonical Wnt signaling activity. Nuclear β-catenin localization provides a more direct readout of signaling dynamics as it tracks the pool of β-catenin available to coactivate Wnt target genes. Live imaging of fluorescently labeled β-catenin allows this nuclear pool to be tracked over time and has been used to study embryonic axis formation, e.g. in sea urchin and *Nematostella* (Weitzel et al. 2004; Lebedeva et al. 2025).

*Hydra* is a small freshwater animal known for its remarkable regenerative capacity. *Hydra’s* body is organized along a single oral-aboral axis, extending from the head at the oral end to the foot at the aboral end. Historically, *Hydra* has served as an important model for studying morphogenesis, contributing to the development of many fundamental concepts in the field (Braun and Keren 2018). The Wnt/β-catenin pathway is widely regarded as the principal signaling pathway involved in establishing the oral-aboral axis in *Hydra* and as a key activator of head formation (Hobmayer et al. 2000; Lengfeld et al. 2009; Nakamura et al. 2011). At the oral end, the head organizer acts as a localized signaling center that maintains the head and patterns the surrounding tissue (Bode 2012). Components of the canonical Wnt/β-catenin pathway, including multiple Wnt ligands, TCF, and β-catenin, are enriched in this region, with β-catenin also exhibiting nuclear localization in cells surrounding the mouth (Hobmayer et al. 2000; Lengfeld et al. 2009; Nakamura et al. 2011). Pharmacological inhibition of GSK3 stabilizes β-catenin, induces its nuclear accumulation throughout the body column, and confers head-organizer activity on body-column tissue (Broun et al. 2005; Nakamura et al. 2011). The mouth at the center of the organizer also coincides with an aster-shaped defect in the alignment of the supracellular ectodermal actin fibers (Maroudas-Sacks et al. 2021, 2025). This defect is essential for mouth function, as contraction of the radially arranged fibers drives the tissue deformation required for mouth opening (Carter et al. 2016).

The Wnt/β-catenin pathway is crucial for axial patterning during head regeneration and budding in *Hydra* (Hobmayer et al. 2000). Grafting experiments show that localized Wnt/β-catenin activity can induce a new head organizer (Broun et al. 2005; Wang et al. 2020), while global Wnt or β-catenin overexpression results in multi-headed animals (Gee et al. 2010; Ferenc et al. 2021). During regeneration from bisected *Hydra*, transcriptional activation of β-catenin and several of its target genes occurs initially at both head- and foot-regenerating wounds (Gufler et al. 2018; Cazet et al. 2021; Tursch et al. 2022). This response persists at the head-regenerating site but is subsequently suppressed at the foot-regenerating site (Gufler et al. 2018; Cazet et al. 2021). How these initially similar wound responses diverge to produce distinct morphogenetic outcomes at the two regenerating ends remains unknown (Gufler et al. 2018; Cazet et al. 2021; Tursch et al. 2022; Wang et al. 2023).

Previous studies in *Hydra* have primarily relied on *in situ* hybridization or transcriptomic analyses at discrete time points to study Wnt/β-catenin signaling (Hobmayer et al. 2000; H. O. Petersen et al. 2015; Gufler et al. 2018; Cazet et al. 2021; Nuninger et al. 2026). Live imaging of a GFP reporter driven by the β-catenin promoter provided additional information about β-catenin expression dynamics, but did not reveal β-catenin protein abundance or its subcellular localization (Iachetta et al. 2018). This distinction is particularly important because of β-catenin’s dual function as a transcriptional coactivator in canonical Wnt signaling and as a structural component of cell–cell junctions. In this work, we use the β-catenin-GFP transgenic strain developed by Nakamura et al. (Nakamura et al. 2011) for high-resolution live imaging, allowing us to obtain a dynamic readout of β-catenin abundance and the signaling-relevant nuclear β-catenin pool throughout regeneration.

We examine β-catenin dynamics *in vivo* in regenerating *Hydra*, across several tissue configurations that differ in their initial geometry, inherited tissue organization, and wound sites. We first examine bisected *Hydra*, the most widely studied regeneration setting, in which transcriptional regulation and proteomic changes have been extensively studied (Hobmayer et al. 2000; Lengfeld et al. 2009; H. O. Petersen et al. 2015; Gufler et al. 2018; Cazet et al. 2021; Nuninger et al. 2026). We then examine tissue rings, which have oral and aboral wound interfaces similar to those in bisected animals, but initially lack both a head and a foot.

Comparing these preparations allows us to examine whether β-catenin dynamics at the two regenerating ends are influenced by signals from pre-existing head or foot structures. Finally, we analyze smaller rectangular tissue fragments, which undergo substantial deformation as they fold into hollow spheroids and have a wound-closure region spanning a large fraction of the tissue (Livshits et al. 2017; Shani-Zerbib et al. 2022; Maroudas-Sacks et al. 2025). Across these configurations, we find that nuclear β-catenin becomes detectable a few hours after excision and is initially enriched across a broad region around the future head-forming site.

Nuclear β-catenin subsequently exhibits non-monotonic dynamics, with the initial broad increase followed by a decrease and later, the emergence of a spatially focused nuclear signal that persists in the developing head. Our measurements also indicate that different initial tissue configurations converge on a localized nuclear β-catenin pattern at the future head through distinct spatiotemporal trajectories.

## Results

To better understand the biochemical signaling processes underlying axis formation during *Hydra* morphogenesis, we follow β-catenin dynamics in live regenerating tissues. We focus on nuclear β-catenin localization because, in the canonical Wnt pathway, β-catenin must enter the nucleus to activate transcription of Wnt target genes (Rim et al. 2022). We utilize a previously developed transgenic *Hydra* strain that expresses β-catenin-GFP in the ectoderm, driven by the endogenous β-catenin promoter (Nakamura et al. 2011). Expression from the endogenous promoter results in reporter levels and localization comparable to those of the endogenous Hyβ-catenin (Nakamura et al. 2011). Although the transgene introduces at least one additional β-catenin copy, it has no obvious morphological consequences. This contrasts with β-catenin overexpression, which increases signaling activity and induces a multiple-head phenotype (Gee et al. 2010), indicating that reporter expression does not substantially perturb β-catenin signaling. The low expression levels, however, generate a relatively low GFP signal. We optimize the imaging conditions to obtain a detectable signal while minimizing photobleaching, using the lowest illumination intensity that provides sufficient signal (Methods). By comparing recordings exposed to different amounts of illumination, we show that the observed dynamics primarily reflect intrinsic changes in the samples rather than photobleaching (Fig. S1).

In mature animals, β-catenin is present at cell–cell junctions throughout the body (Fig. 1A,B). β-catenin also localizes to the nuclei of cells surrounding the mouth, where it mediates canonical Wnt signaling (Fig. 1A,B) (Nakamura et al. 2011). In the tentacles, we observe prominent β-catenin-positive plaques at the basal ectodermal surface (Fig. 1B), likely associated with the specialized junctional complexes that anchor nematocytes to ectodermal battery cells (Wood and Novak 1982; Campbell 1987). Treatment with Alsterpaullone, which inhibits GSK3 thereby stabilizing β-catenin, generates an increase in β-catenin-GFP levels and induces widespread nuclear localization (Fig. S2), consistent with previous reports (Broun et al. 2005; Nakamura et al. 2011). The live β-catenin-GFP reporter thus reproduces both the established β-catenin localization pattern in mature animals and the response to pharmacological β-catenin stabilization, providing a reliable readout of changes in both β-catenin abundance and nuclear localization (Nakamura et al. 2011).

**Figure 1.**
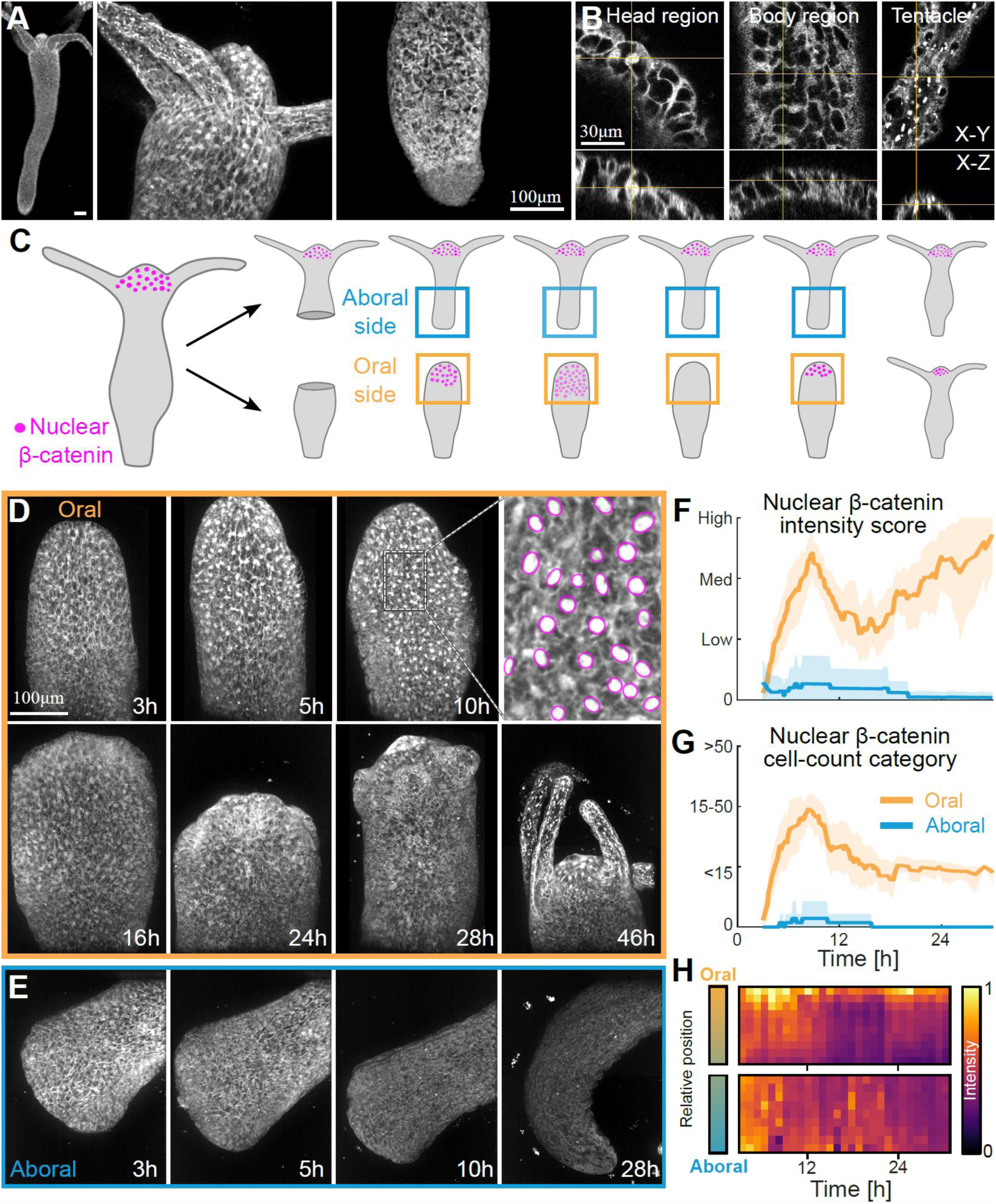
Live imaging of β-catenin dynamics during head and foot regeneration in bisected Hydra. (A) Airyscan confocal projected image of a mature Hydra expressing β-catenin-GFP in the ectoderm (left) with magnified views of the head and foot (right). (B) Airyscan confocal images showing x-y (top) and x-z (bottom) cross-sections through regions of the head, body column, and tentacle. The images show β-catenin localization at cell-cell junctions throughout the animal, nuclear localization in the head region (left), and localization in basal plaques in the tentacles (right). (C) Schematic illustration of the regeneration of bisected Hydra and the dynamics of nuclear β-catenin localization (magenta) during this process. (D,E) Spinning-disk confocal projected images from time-lapse recordings of the head-regenerating oral end (D) and foot-regenerating aboral end (E) of bisected Hydra expressing β-catenin-GFP in the ectoderm (Movies 1 and 2, respectively). A magnified view depicts nuclei exhibiting detectable nuclear β-catenin localization (magenta; D, top right panel). Time from bisection is indicated. (F,G) Time course of nuclear β-catenin localization in bisected animals during head (orange) and foot (cyan) regeneration. Imaging started 3-5 h after bisection, following sample preparation and microscope mounting. Nuclear β-catenin intensity (F) and the number of nuclei exhibiting detectable nuclear β-catenin localization (G) were manually scored at each time point in N=16 regenerating head samples and N=15 regenerating foot samples. Intensity was scored as none, low, intermediate, and high (Fig. S4; Methods). Estimated cell counts were assigned to four categories: 0 (no nuclei), <15 nuclei, 15-50 nuclei, >50 nuclei. Lines indicate the mean score, and shaded regions indicate the 95% bootstrap confidence interval. (H) Kymographs showing total β-catenin-GFP fluorescence intensity along the normalized oral-aboral axis (y-axis) over time (x-axis) within the oral half of the head-regenerating piece (top) and the aboral half of the foot-regenerating piece (bottom), for the bisected animals shown in (D) and (E), respectively (Methods).

We first examine β-catenin dynamics during head and foot regeneration in bisected animals. During the first hours after bisection, nuclear β-catenin localization is not detectable at either the oral or aboral-facing wound sites, which respectively correspond to the future head and foot sites (Fig. 1C-G). Several hours later, a pronounced asymmetry in nuclear β-catenin localization emerges between the oral and aboral sides: a broad region of cells exhibiting nuclear β-catenin becomes apparent at the future head side, whereas little or no nuclear localization is detectable at the future foot side (Fig. 1D-E; Movies 1, 2). Kymographs of total β-catenin-GFP fluorescence suggest an early increase in β-catenin abundance at both regenerating ends (Fig. 1H), even though nuclear localization is detected only at the future head site. Subsequently, nuclear β-catenin in the future head region exhibits non-monotonic dynamics: following the initial broad increase in nuclear signal ∼4-8 h after bisection, the signal decreases before increasing again at ∼18 h within a smaller region near the tip of the regenerating head (Fig. 1F,G; Movie 1). This localized signal persists and develops into the characteristic stable nuclear β-catenin pattern observed in cells surrounding the mouth of mature *Hydra*. In contrast, nuclear β-catenin typically remains undetectable at the foot-regenerating end throughout regeneration.

We next examine β-catenin dynamics in regenerating tissue rings, in which the head and foot are both removed (Fig. 2). Tissue rings retain their original oral-aboral axis, which becomes the body axis of the regenerated animal, with a head forming at the site of the oral-facing wound and a foot at the aboral-facing wound (Javois et al. 1986; Livshits et al. 2022). The β-catenin dynamics at the oral- and aboral-facing wound sites within a regenerating tissue ring are similar to those observed at the respective wound sites in bisected animals (Fig. 2B,E).

**Figure 2.**
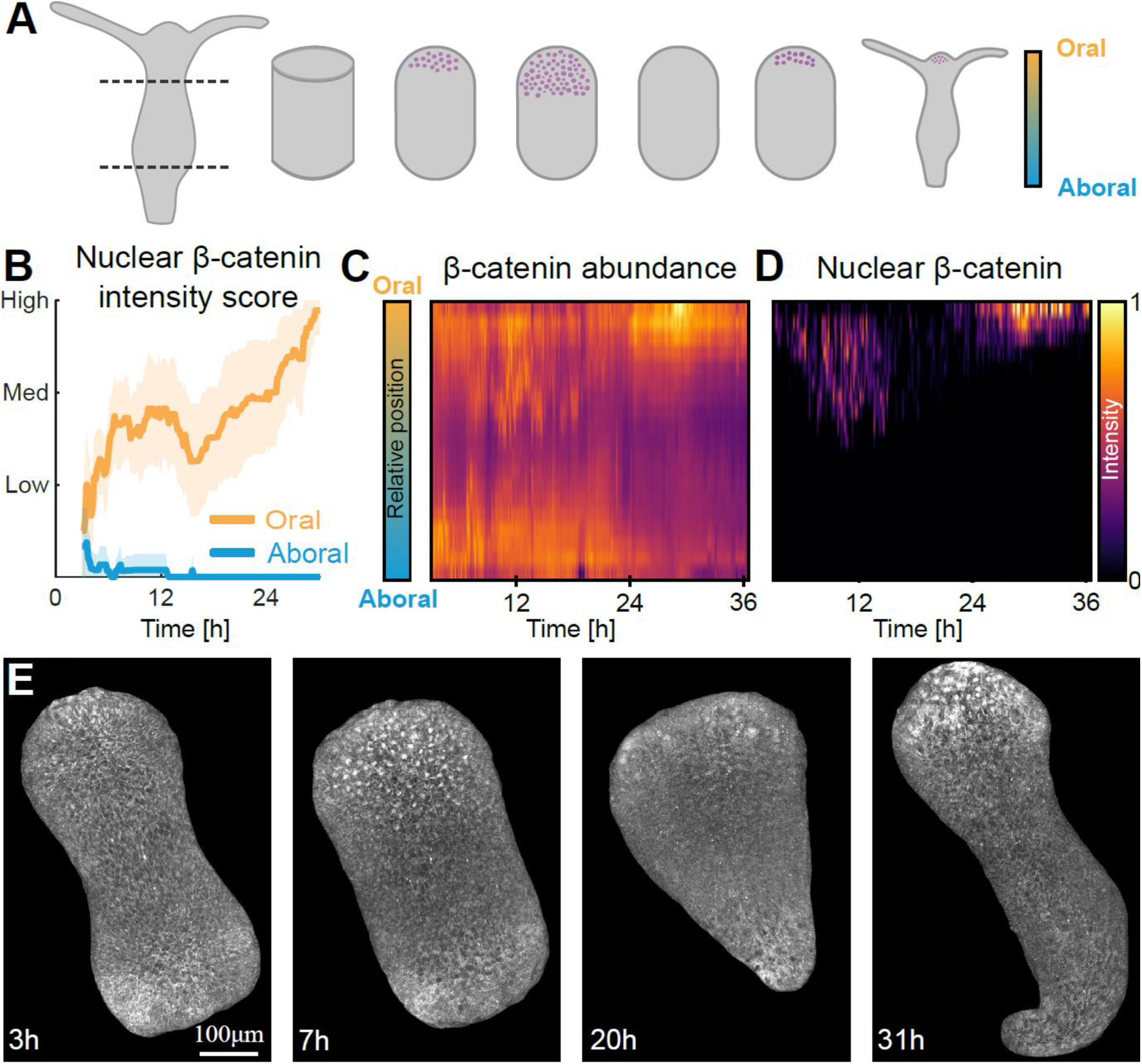
Live imaging of β-catenin dynamics in regenerating Hydra tissue rings. (A) Schematic illustration of the regeneration of an excised tissue ring and the dynamics of nuclear β-catenin localization (magenta) during this process. (B) Time course of nuclear β-catenin localization at the oral (orange) and aboral (cyan) sides of regenerating tissue rings (N=18). Imaging started 3-4 h after excision, following sample preparation and microscope mounting. Nuclear β-catenin-GFP intensity was manually scored as in Fig. 1F (Methods). Lines indicate the mean score, and shaded regions indicate the 95% bootstrap confidence interval. (C,D) Kymographs showing the total β-catenin-GFP fluorescence intensity (C) and nuclear β-catenin-GFP localization (D) along the normalized oral-aboral axis (y-axis) over time (x-axis) for the tissue ring shown in E (see Methods for details). (E) Spinning-disk confocal projected images from a time-lapse movie of a regenerating ring expressing β-catenin-GFP (gray; Movie 3). Time from excision is indicated.

During the first hours after excision, nuclear β-catenin localization is not detectable at either side. After several hours, nuclear localization becomes strongly asymmetric: nuclear β-catenin is observed across a broad region on the originally oral-facing side, whereas little or no nuclear localization is detected on the originally aboral-facing side (Fig. 2; Movie 3). The subsequent spatiotemporal patterns of β-catenin nuclear localization on the oral- and aboral-facing sides further resemble those on the future head and foot sides of bisected animals, respectively. On the oral-facing side, the initial broad signal decreases before reemerging in a spatially focused region at the tip, while on the aboral-facing side, nuclear localization remains weak or undetectable (Fig. 2B). Kymographs along the regeneration axis provide a continuous view of the changes in total β-catenin levels and nuclear localization, and the emergence of asymmetry between the future head and foot sites (Fig. 2C,D). The development of this asymmetry within a regenerating tissue ring that lacks both a functional head and foot indicates that existing terminal structures are not required for this polar β-catenin localization.

Finally, we examine β-catenin dynamics in regenerating tissue fragments (Fig. 3). The smaller size of tissue fragments compared with bisected animals and tissue rings makes it possible to use Airyscan confocal imaging despite its slower acquisition speed. This provides higher spatial resolution and signal-to-noise ratio than spinning-disk imaging, enabling automated identification of nuclei with nuclear β-catenin localization (Methods). As an excised tissue fragment folds and closes into a hollow spheroid, its initially head-facing edge comes into contact with and attaches to its initially foot-facing edge (Fig. 3A) (Livshits et al. 2017; Shani-Zerbib et al. 2022; Maroudas-Sacks et al. 2025). Imaging during this early folding step shows that nuclear β-catenin localization is not detected immediately following excision, but emerges only after the fragment has folded into a spheroid (Fig. S3). Despite the extensive tissue deformation involved in this process, nuclear β-catenin dynamics in regenerating fragments share key features with those observed in bisected animals and tissue rings. In all cases, nuclear β-catenin localization first arises in a broad region, exhibits non-monotonic dynamics, and eventually stabilizes in a small region around the future head site (Fig. 3B,C,F; Movie 4). Nevertheless, important differences emerge in both the spatial distribution and the temporal dynamics of nuclear β-catenin localization.

**Figure 3.**
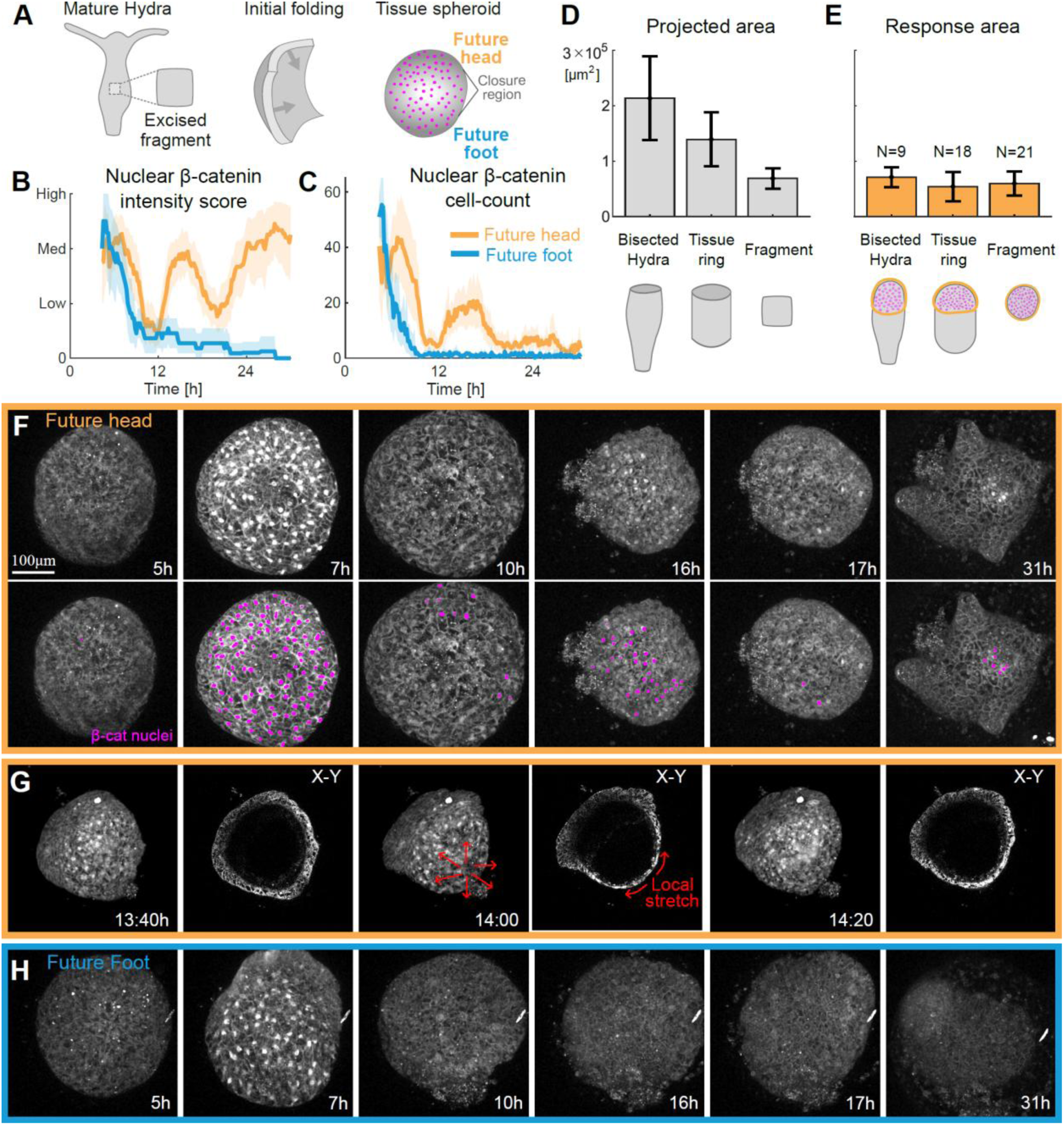
Live imaging of β-catenin dynamics in regenerating Hydra tissue fragments. (A) Schematic illustration of an excised tissue fragment, its initial folding, and the broad distribution of nuclear β-catenin (magenta) in the tissue spheroid during the initial peak. (B, C) Nuclear β-catenin intensity score (B) and number of nuclei exhibiting detectable nuclear β-catenin localization (C) as a function of time in the future head (orange; N=21) and future foot (cyan; N=11) regions. Regenerating Hydra tissue fragments expressing β-catenin-GFP in the ectoderm were followed throughout regeneration. Imaging started 3-5 h after excision, following sample preparation and microscope mounting. The future head- and foot-forming regions in spheroids were identified by tracking backward through the time-lapse recordings (Methods). Analysis is based on recordings in which either the future head or the future foot region remained predominantly facing the objective throughout the regeneration process. Nuclei exhibiting detectable nuclear β-catenin localization were automatically identified (Fig. S5), and the intensity score was manually assigned as in Fig. 1F (Methods). Lines indicate the mean, and shaded regions indicate the 95% bootstrap confidence interval. (D) Bar plot of the projected tissue area of bisected animals, tissue rings, and tissue fragments (Methods). (E) Bar plot of the maximal response area in the same samples, defined as the projected tissue area containing nuclei with detectable nuclear β-catenin localization when the initial response is broadest (Methods). In D,E, bars indicate the mean and error bars indicate the standard deviation. (F) Projected images from an Airyscan confocal time-lapse movie of a regenerating tissue fragment expressing β-catenin-GFP (gray) showing the future head region (Movie 4). Automatically detected nuclei with nuclear β-catenin localization are overlaid in magenta in the lower panels (Methods). The initial peak of nuclear β-catenin localization is widespread, whereas the subsequent increase in nuclear β-catenin levels is more localized to the future head region. (G) Images from an Airyscan confocal time-lapse movie of a regenerating tissue fragment expressing β-catenin-GFP at three time points before (left), during (center) and after (right) a transient global deformation event involving localized tissue stretching. For each time point, a projected view of β-catenin localization (left) is shown alongside a cross-section highlighting tissue stretching (right). The region of nuclear β-catenin localization is centered at the site of localized tissue stretching. (H) Projected images from an Airyscan confocal time-lapse movie of a regenerating tissue fragment expressing β-catenin-GFP (gray) showing the future foot region (Movie 6).

The initial peak of nuclear β-catenin localization in fragments occupies a large portion of the regenerating tissue, often encompassing nearly the entire spheroid (Fig. 3F,H; Movies 4, 5). In contrast, the corresponding response in bisected animals and tissue rings occupies only a fraction of the regenerating tissue (Figs. 1,2). Thus, while the initial response extends over regions of comparable size across all sample types, the smaller overall size of fragments means that this response often encompasses both the future head and foot regions (Fig. 3D-F,H). The transient presence of nuclear β-catenin in the future foot region in regenerating fragments also indicates that this early nuclear response alone is not sufficient to specify head formation.

A further distinction is observed in the temporal dynamics of nuclear β-catenin localization, which are more complex in fragments than in bisected animals and tissue rings. Following the initial peak at the oral-facing side, nuclear β-catenin typically decreases but remains detectable in bisected animals and tissue rings (Figs. 1,2). In fragments, in contrast, nuclear β-catenin at the future head region typically falls below detectable levels before reappearing in an additional peak, declining again, and only then becoming stably localized within a small region surrounding the future head site (Figs. 3B,C,F, S5). Thus, the temporal profile in fragments typically contains two distinct minima (“valleys”) between successive periods of nuclear β-catenin localization, whereas bisected animals and tissue rings exhibit only one.

Spatially, the subsequent peaks of nuclear β-catenin localization, following the initial nearly system-wide response, are centered in the region undergoing recurring localized tissue stretching (Fig. 3G). In previous work, we showed that such recurring mechanical strain focusing is characteristic of regenerating *Hydra* and occurs at the location of a defect in the nematic alignment of the supracellular actin fibers at the future head site (Maroudas-Sacks et al. 2025). During global tissue contractions, the radial arrangement of the contracting actomyosin fibers focuses strain at the defect site, causing cells at the defect core to stretch to more than twice their initial area. In cross-section, this localized stretching is evident as pronounced thinning of the ectoderm (Fig. 3G). The β-catenin-GFP signal at cell–cell junctions provides additional information about changes in cell shape during these stretching events (Movie 4). Together, these features allow us to identify the strain-focusing region associated with the future head defect and show that it colocalizes with the center of the nuclear β-catenin localization peak, even without actin labeling. Although head formation remains broadly biased toward the oral-facing side of the fragment, the initial tissue folding brings the original oral- and aboral-facing edges into contact (Fig. S3), and the future head site is not located at the fragment’s original oral-facing edge (Shani-Zerbib et al. 2022; Maroudas-Sacks et al. 2025).

Altogether, our results reveal common features of nuclear β-catenin localization across different regeneration geometries, alongside important differences in their dynamics. In all cases, an initially broad nuclear β-catenin response is followed by localization to a much smaller region at the future head site, but the trajectory toward this shared endpoint varies with the size and initial configuration of the regenerating tissue. Across the different geometries, the localized nuclear β-catenin peaks are centered around the actin-fiber defect sites, which experience recurring strain focusing.

## Discussion

Wnt/β-catenin signaling has a central role in axial patterning and head-organizer formation in *Hydra*, yet the mechanisms through which the processes underlying axial patterning are spatiotemporally organized during regeneration to robustly establish the body axis remain poorly understood. One important aspect of axial patterning is the transition from an initial wound response to distinct regeneration trajectories. Previous transcriptional studies indicate that the early injury-induced gene expression response in bisected animals is largely shared between head- and foot-regenerating wounds, with distinct head- and foot-specific transcriptional patterns emerging approximately 6–12 h after excision (Gufler et al. 2018; Cazet et al. 2021). Our live measurements reveal a corresponding transition at the level of β-catenin protein localization (Fig. 1). We find that pronounced nuclear localization emerges at approximately 4-8 h specifically at the oral-facing end of bisected animals and tissue rings (Figs. 1F,2B). The timing and spatial localization of this initial nuclear β-catenin peak therefore correlate with the transition from a generic wound response to distinct head- and foot-specific transcriptional patterns. Interestingly, the transient presence of nuclear β-catenin in the future foot region in small regenerating fragments demonstrates that early nuclear localization alone is insufficient to specify head identity (Fig. 3B).

The increase in total β-catenin abundance at the foot-regenerating end may reflect a general response to injury rather than sustained activation of canonical Wnt signaling. Early *Hydra* regeneration is associated with broad protein stabilization and increased transcription (H. O. Petersen et al. 2015), and Wnt pathway components, including β-catenin, are initially transcriptionally upregulated following both head and foot amputation (Gufler et al. 2018; Cazet et al. 2021). Thus, reduced protein degradation and/or increased β-catenin synthesis could contribute to the early rise in β-catenin at both wounds. At the aboral-facing wound, this increase in β-catenin abundance is not accompanied by detectable nuclear accumulation. However, we cannot exclude a weak nuclear β-catenin response below our detection limit, and therefore cannot determine whether the early injury-induced Wnt response during foot regeneration is β-catenin-independent. The absence of a pronounced nuclear increase during foot regeneration nevertheless indicates that β-catenin abundance alone does not determine its nuclear localization.

Our comparison of bisected animals and tissue rings further constrains what determines the divergence of regeneration toward head or foot formation. In bisected animals, the regenerating tissue remains connected to an existing head or foot, which could potentially provide long-range signals that influence regeneration at the wound. Tissue rings, in contrast, initially contain neither structure. Nevertheless, we find that rings exhibit strikingly similar β-catenin dynamics, with nuclear localization emerging only at the oral-facing side and subsequently becoming focused at the future head site (Figs. 1,2). Thus, although long-range signals may contribute to regeneration, signals originating from an existing head or foot are not required for the emergence of the observed asymmetry in nuclear β-catenin localization or for the distinct head- and foot-regeneration trajectories. Instead, the ability of isolated rings to reproduce the dynamics observed following bisection indicates that sufficient positional information is retained within the excised body-column tissue itself.

An important feature of the nuclear β-catenin response is its non-monotonic temporal dynamics. Previous measurements of β-catenin transcript levels during head regeneration showed an early increase followed by a decrease and a subsequent increase at later stages (Lommel et al. 2018; Nuninger et al. 2026). Recent time-resolved transcriptional analysis further revealed dynamic changes in Wnt ligand expression and canonical Wnt pathway activity during head regeneration, with diminished canonical Wnt-associated gene expression and reduced nuclear β-catenin levels observed during an intermediate period (Nuninger et al. 2026). Here, we characterize the non-monotonic nuclear β-catenin dynamics in detail in live regenerating Hydra across different initial conditions. In bisected animals, an initial peak in nuclear β-catenin is followed by a decline and subsequent re-emergence before stable localization at the future head is established. Following the same regenerating tissue by live microscopy allows these transient changes to be resolved at high temporal resolution. This is particularly important in fragments, where the temporal profile typically contains two distinct minima in nuclear β-catenin levels, compared with a single minimum in bisected animals and tissue rings. Despite these differences in temporal dynamics, the different initial tissue configurations robustly converge on the same outcome: stable nuclear β-catenin localization at the future head. The variation in the trajectories across these different tissue contexts suggests that β-catenin signaling is influenced by additional inputs during organizer establishment, rather than following a fixed temporal program.

The initial nuclear β-catenin response extended over regions of similar size across the different sample types, despite differences in their overall size (Fig. 3D,E). In bisected animals and tissue rings, the response was confined to the oral-facing end and subsequently narrowed toward the future head site. In small fragments, however, the initial response extended over nearly the entire tissue before becoming restricted to a small region at the future head site. Thus, in bisected animals and rings, the initial response already distinguishes the head-forming end, whereas in fragments it does not. This raises the question of what provides the positional information that specifies the head-forming site within such a broad initial response region.

A possible source of such positional information is the supracellular actin-fiber organization and its associated pattern of topological nematic defects. Our previous work showed that the future head forms at an aster-shaped defect in the alignment of the supracellular actin fibers across different regeneration geometries (Maroudas-Sacks et al. 2021, 2025). This association is particularly striking in regenerating fragments, where the initial folding transforms the planar fragment into a closed, hollow spheroid, bringing the originally oral- and aboral-facing edges into contact (Fig. S3). Although tissue polarity is largely retained, this rearrangement makes the spatial distinction between the oral and aboral regions less straightforward than in bisected animals and tissue rings, and the new head does not simply emerge from the original head-facing edge (Shani-Zerbib et al. 2022; Maroudas-Sacks et al. 2025). Here we show that following the initial nearly system-wide response, the localized nuclear β-catenin peaks in regenerating fragments are centered on the region exhibiting the characteristic stretching events (Fig. 3G), previously shown to coincide with the actin-fiber defect (Maroudas-Sacks et al. 2025). This spatial correspondence is particularly interesting given β-catenin’s dual roles as a component of cell–cell adhesion complexes and as a nuclear effector of canonical Wnt signaling, making it a potential mediator of coupling between tissue mechanics and biochemical signaling (Fagotto 2013). This raises the possibility that mechanical deformation at the defect site could influence β-catenin localization and dynamics. For example, localized tissue stretching at defect sites could alter the association or turnover of β-catenin at cell–cell junctions or modulate β-catenin nuclear translocation. At present, however, our evidence for such coupling is limited to the spatial correlation. Future work will be required to determine whether and how mechanical inputs influence β-catenin dynamics and, more generally, the signaling processes associated with head-organizer formation.

Our results demonstrate the value of following β-catenin protein localization continuously in live *Hydra*. Protein dynamics cannot be inferred directly from the more commonly studied transcriptional responses, because protein abundance and localization are subject to additional layers of regulation. This is particularly relevant during early regeneration, when the wound response involves rapid changes in protein phosphorylation and stability (H. O. Petersen et al. 2015; Tursch et al. 2022), and changes in protein abundance substantially differ from transcript level changes (H. O. Petersen et al. 2015). Moreover, subcellular-resolution imaging reveals the distribution of β-catenin among its junctional, cytoplasmic, and signaling-relevant nuclear pools within the same regenerating animal over time. Our live imaging of β-catenin–GFP in regenerating *Hydra* thus complements previous studies of Wnt/β-catenin signaling by providing a direct readout of nuclear β-catenin localization at high temporal resolution.

Together, our results show that the spatiotemporal dynamics of nuclear β-catenin are non-monotonic and vary with the initial tissue configuration, yet converge on a shared endpoint characterized by spatially restricted nuclear β-catenin at the future head. Across the different configurations, nuclear β-catenin progresses from an initially broad distribution toward a highly focused final pattern. In regenerating fragments, an additional peak of intermediate spatial extent is observed during the intervening period. Our observations further reveal a spatial correlation between nuclear β-catenin localization and the site of a nematic defect in the organization of the supracellular actin fibers. Future work combining live β-catenin imaging with controlled perturbations of Wnt signaling, tissue mechanics, and cytoskeletal dynamics will be required to determine what establishes the spatiotemporal pattern of nuclear β-catenin localization and whether mechanical inputs contribute to its formation. Resolving the processes responsible for the repeated increases and decreases in nuclear β-catenin and changes in its spatial extent will be important for understanding how different dynamic signaling trajectories converge on robust axial patterning.

## Methods

### Hydra strains and culturing

All experiments are performed with a transgenic strain of *Hydra* vulgaris (AEP) expressing EGFP-tagged Hyβ-catenin in the ectoderm (Nakamura *et al*. 2011) developed by the Holstein lab (University of Heidelberg), and generously provided by Charisios Tsiairis (Friedrich Miescher Institute for Biomedical Research, Basel, Switzerland). Animals are cultivated in *Hydra* medium (HM; 1mM NaHCO3, 1mM CaCl2, 0.1mM MgCl2, 0.1mM KCl, and 1mM Tris-HCl, pH7.7) at 18°*C*. The animals are fed with live Artemia nauplii three times a week and subsequently washed ∼6-8 hours after feeding. Experiments are done ∼24 hours after feeding.

### Sample preparation

Different tissue segments are excised from mature *Hydra* using a scalpel equipped with a #15 blade as follows. Bisected animals are formed by making a transverse cut in the middle of a mature *Hydra*. Tissue rings (comprising ∼½ of the original animal’s length) are excised by removing the head and foot regions of a mature *Hydra*. Rectangular tissue fragments are formed by first excising two tissue rings (each comprising ∼¼ of the original animal’s length) from the gastric region of a mature animal, and then further cutting each ring into 2-4 parts by additional longitudinal cuts.

Pharmacological enhancement of the Wnt/β-catenin signaling pathway is done with Alsterpaullone (ALP; Sigma) by incubating mature *Hydra* in 5µM ALP in HM for 24 hours.

For regeneration experiments, excised tissues are first allowed to seal in HM for ∼2-3 h. Subsequently, the samples are immersed in liquefied 0.5% clear low gelling agarose (sigma) prepared with HM, that has sufficiently cooled (∼35°C) and placed in preformed wells, made from 2% agarose (Sigma) prepared with HM in 50 mm glass-bottom Petri dishes (Fluorodish). After the 0.5% gel solidifies, 3-4 ml HM are added above the sample. The 0.5% low gelling agarose gel around the samples reduces tissue movement during imaging, allowing us to find samples that remain sufficiently stable to follow the same tissue regions over time.

To visualize full animals, animals are relaxed in 2% urethane in HM and placed between two coverslips separated by ∼200 µm using a double-sided tape spacer.

### Microscopy

3D time-lapse movies of regenerating *Hydra* are acquired at room temperature in one of two microscopy systems. The first is an upright spinning-disk confocal microscope equipped with a Yokogawa CSU-X1 spinning-disk unit (Intelligent Imaging Innovations, Slidebook software). The EGFP-tagged Hyβ-catenin is excited using a 200 mW 488 nm laser. Time-lapse movies are acquired with a sCMOS camera with 2×2 or 4×4 binning (Andor Zyla 4.1) using a 20× dipping objective (NA=0.5) from above. Since imaging from above requires an open dish, the Petri dish is equipped with a homemade Teflon ring with tubing connected to a peristaltic pump (Ismatec), to allow perfusion of media.

The second system is an inverted LSM 980 confocal microscope equipped with an Airyscan2 detector (Zeiss, ZEN software), providing improved image resolution and contrast at the expense of longer acquisition times. The sample is excited using a 30 mW 488 nm laser at 10% laser power, through a 25× glycerol objective (NA=0.8).

In both systems, 3D time-lapse movies are acquired with a z-interval of 3-6 μm, and a time interval ranging from 5 minutes to 1 hour, over 2-3 days, until regeneration is complete. The 3D stacks span the side of the sample facing the objective over a range of 110-150 μm.

For all sample types, only the tissue facing the objective can be visualized, as image degradation prevents visualization of the opposite side. Tissue rings and bisected animals typically maintain a side-view orientation, with the future body axis approximately parallel to the imaging plane. In contrast, spheroids formed from small tissue fragments can adopt arbitrary orientations. Embedding in gel reduces rotation during imaging, allowing the future head- and foot-forming regions in spheroids to be identified retrospectively by tracking these regions backward through the time-lapse recordings. For analysis, we selected recordings in which either the future head or the future foot region remained predominantly facing the objective during the regeneration process.

The fluorescence signal is typically very low since β-catenin is expressed from its endogenous promoter. We used the lowest excitation intensities that still allowed the fluorescence dynamics during regeneration to be reliably followed. The imaging modality is selected to obtain sufficient sensitivity for the required field of view for each type of sample. Spheroids formed from small, excised tissue fragments are imaged using Airyscan microscopy, which provides improved spatial resolution and contrast, and enables us to identify β-catenin nuclear localization using automated analysis (see below). Tissue rings are imaged using the spinning-disk confocal system. As most rings are too large to fit within a single field of view, images are acquired at several partially overlapping positions and subsequently stitched as described below. The spinning-disk system is used for this multi-position acquisition because its faster imaging speed allows all positions to be acquired before substantial sample movement or deformation occurs, enabling acceptable image stitching.

Bisected animals are relatively large and exhibit more pronounced movement than tissue rings or fragments during regeneration. Consequently, imaging the entire sample using fixed multi-position acquisition would require a large number of adjacent fields of view, making acquisition impractically slow. We therefore image only the regenerating head- or foot-forming end, which is either followed manually, typically at 1-h intervals, by repositioning the sample before each acquisition, or imaged automatically at fixed positions. In the latter case, multiple samples are imaged in parallel, and recordings are subsequently selected for analysis if the regenerating end remains within the field of view for a sufficient fraction of frames to follow its dynamics. Bisected animals were imaged using either spinning-disk confocal microscopy or Airyscan microscopy. For measurements of total tissue size, overview images were acquired using a separate 5× objective positioned below the sample in the spinning-disk system’s up-and-under configuration.

Full animals are imaged in the inverted LSM 980 confocal microscope using a 10× (NA=0.5) air objective or a 25× (NA=0.8) glycerol objective with a z-interval of 1.5-3 μm.

### Image processing and analysis

To characterize nuclear β-catenin abundance and localization, we use 2D maximum-intensity projections of the acquired 3D image stacks (acquired at 3-6 μm z-intervals), which facilitate the identification and analysis of nucleus-shaped objects in the sample. For samples imaged at multiple positions, image stitching is performed as follows. The 2D projections taken from several adjacent images with an overlap (∼ 50-100 μm) are stitched using the Pairwise Stitching plugin in ImageJ (plugin internal version 1.2) (Preibisch et al. 2009). Alignment is computed using the built-in phase correlation algorithm, allowing translational shifts only.

Fused images are generated using maximum intensity fusion as previously described (Westfried et al. 2026). The 2D projections are median-filtered in ImageJ with a radius of 1.25 μm to reduce salt-and-pepper noise, and assembled into time-lapse movies to visualize β-catenin dynamics over time.

To compare the size of the different types of samples and their maximal response area, we used projected areas. For each movie, we first manually selected the time point at which nuclei with detectable nuclear β-catenin localization were distributed over the largest tissue area during the initial response peak. At this selected time point, the projected sample area was determined by segmenting the entire tissue based on the fluorescence signal and measuring its projected area (Fig. 3D). The region with nuclear β-catenin localization was manually outlined on the projected image based on the detected β-catenin-positive nuclei, and its area was measured to obtain the maximal response area (Fig. 3E).

Mean time courses of nuclear β-catenin intensity scores (Figs. 1F, 2B, 3B, and S1B), nuclear count categories (Fig. 1G), and nuclear counts (Fig. 3C) were calculated by averaging across samples at each time point, omitting missing values. Pointwise 95% confidence intervals were estimated using 1,000 bootstrap resamples. Each resample contained the same number of samples as the original dataset, drawn with replacement. Entire sample trajectories were resampled together, preserving temporal correlations within each recording. The mean was calculated at each time point for each resample, omitting missing values, and confidence limits were defined by the 2.5th and 97.5th percentiles of the bootstrap means.

To generate spatiotemporal β-catenin fluorescence profiles along the body axis of regenerating tissue rings and bisected *Hydra* (Fig. 1H, 2C), we analyze samples that remain positioned on their side throughout the regeneration process. The head- and foot-forming sides are identified from the final frames of the time-lapse movie, when the regenerated head and foot can be distinguished morphologically and tentacles are clearly visible. Their positions are then traced backward frame by frame to the beginning of the movie. The regeneration axis is defined as the central line connecting the head- and foot-forming sides using the *centerline* function in the *Celltool* software package (Pincus and Theriot 2007). For each projected image, the tissue region is segmented using Li thresholding in ImageJ to define a tissue mask. 20 equidistant points are placed along the defined regeneration axis, and the tissue mask is divided into swaths corresponding to the Voronoi regions between adjacent points. For each swath, the mean projected image intensity is calculated. Kymographs of the β-catenin fluorescence signal over time are generated by depicting the mean fluorescence intensity along the regeneration axis, where the distance between the oral and aboral ends has been normalized (Fig. 1H, 2C). For display, intensities were normalized by a fixed multiple of the mean intensity over the corresponding movie. The same multiplicative factor was used for both kymographs in Fig. 1H. Normalization was applied uniformly across all time points within each movie.

To generate kymographs of nuclear β-catenin (Fig. 2D), nuclei are manually annotated (Movie 3). Nuclear masks are generated by placing circular objects with a diameter of 9 pixels at the manually determined nuclei positions. The kymographs along the oral-aboral axis of nuclear β-catenin localization over time are generated by depicting the mean nuclear mask intensity along the regeneration axis as above. The intensity is scaled to half the maximum intensity in the kymograph, saturating < 0.25% of the pixels in the kymograph to better visualize the early dynamics.

Nuclear β-catenin localization is assessed manually from maximum-intensity projections of time-lapse image stacks of regenerating *Hydra*. Nucleus-shaped objects containing a visible β-catenin signal are identified by eye in each projected image, and nuclear localization is scored according to their number and their fluorescence intensity. For each frame, the number of cells with detectable nuclear β-catenin localization is estimated manually and assigned to one of four categories: 0, no visible nuclei; fewer than ∼15 nuclei; ∼15–50 nuclei; and more than ∼50 nuclei (Fig. 1G). Nuclear β-catenin intensity is manually scored from 0 to 3: 0, no detectable nuclear fluorescence; 1, low nuclear fluorescence; 2, intermediate nuclear fluorescence; and 3, high nuclear fluorescence, as illustrated in Fig. S4 (Figs. 1F, 2B, 3B).

In regenerating spheroids from small excised tissue fragments imaged using Airyscan microscopy, nuclear β-catenin localization is also detected automatically using Cellpose, a deep-learning-based segmentation algorithm for biological images implemented in Python (Stringer et al. 2021; Pachitariu et al. 2025). We use the CellPose Segment Anything Model (CPSAM) module, to identify bright nuclear-sized objects in the median-filtered maximum-projection images (Fig. S5). The following parameters are used: pixel size = 0.83 µm, diameter = 20 pixels, cellprob_threshold = 2.0, min_size = 49 pixels (min nuclei size in pixel), max_size_fraction =20^2^/488^2^ (max size relative to image size). To minimize false positive detections of dark objects with bright surroundings, segmented regions are removed if their mean intensity is not sufficiently higher than that of their immediate surroundings.

The surrounding intensity is calculated by dilating the segmented area using a 5 pixel radius and subtracting the segmented regions, so that only a ring remains. The mean intensity of the surrounding ring I_ring_ and the mean intensity of the original segmented region I_seg_ and their ratio 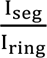 are calculated. A segmented region is kept only if 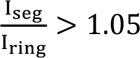 . The number of cells with detectable nuclear β-catenin is determined based on this automated segmentation of nuclei. At each time point, the cell count is determined by counting all identified nuclei (see Fig. 3C). The automated and manual approaches yield comparable results (Fig. S5).

## Acknowledgments

This work was supported by a grant from the Israel Science Foundation to K.K. (grant No. 3565/24). We thank Liron Stettiner for help with automated image analysis, and Nitzan Dahan from the LS&E Imaging Center for advice and help with Airyscan imaging. We thank all the members of our lab for help, discussions and comments on the manuscript. We thank Marko Popovic, Erez Braun, Suat Ozbek, Haguy Wolfenson, Celina Juliano and Jana Fuhrmann for discussions and comments on the manuscript.

## Supplementary Figures

**Figure S1.**
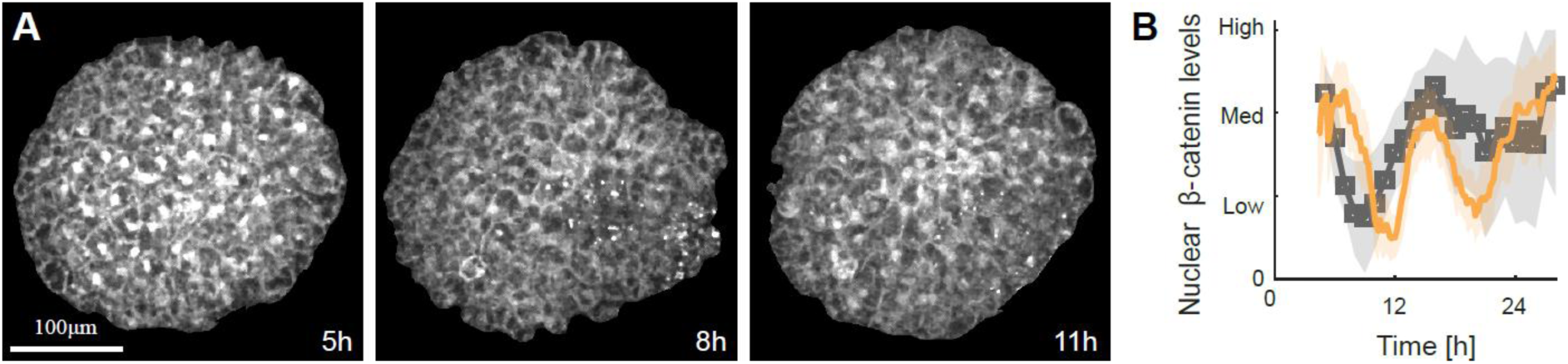
Nuclear β-catenin dynamics at different imaging intervals. We assessed the contribution of photobleaching to the observed dynamics by comparing recordings of samples imaged at time intervals ranging from 5 min to 1 h and therefore exposed to different amounts of illumination. (A) Airyscan confocal projected images from a time-lapse recording of a regenerating tissue spheroid expressing β-catenin-GFP, acquired at 1-h intervals. Three representative time points are shown, showing the initial broad peak (left), subsequent decline (center) and second narrower peak (right). (B) Time evolution of nuclear β-catenin intensity levels, determined manually (Methods), for time-lapse recordings acquired at 5–10- min intervals (orange) or 1-h intervals (gray). The comparable dynamics observed for these conditions indicate that photobleaching does not substantially contribute to the measured signal changes and that the observed dynamics primarily reflect changes in the sample.

**Figure S2.**
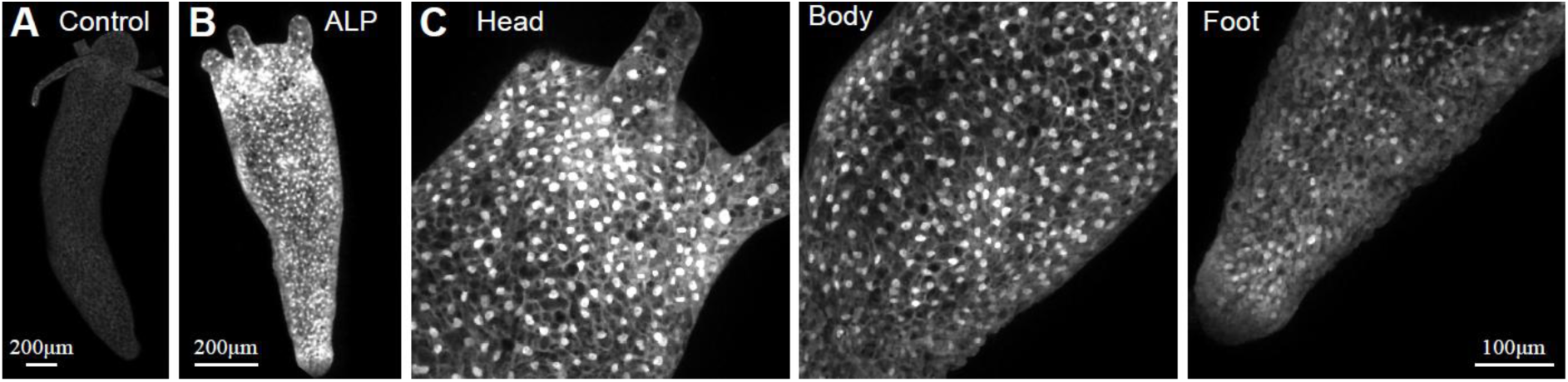
Alsterpaullone (ALP) treatment induces increased β-catenin abundance and widespread nuclear localization. (A,B) Airyscan confocal projected images of control (A) and ALP-treated (B) animals expressing β-catenin-GFP. The images are displayed with identical brightness and contrast settings, highlighting the increased β-catenin abundance following ALP treatment, which stabilizes β-catenin by reducing its degradation. (C) Magnified views of different regions show widespread nuclear β-catenin localization throughout the body following ALP treatment.

**Figure S3.**
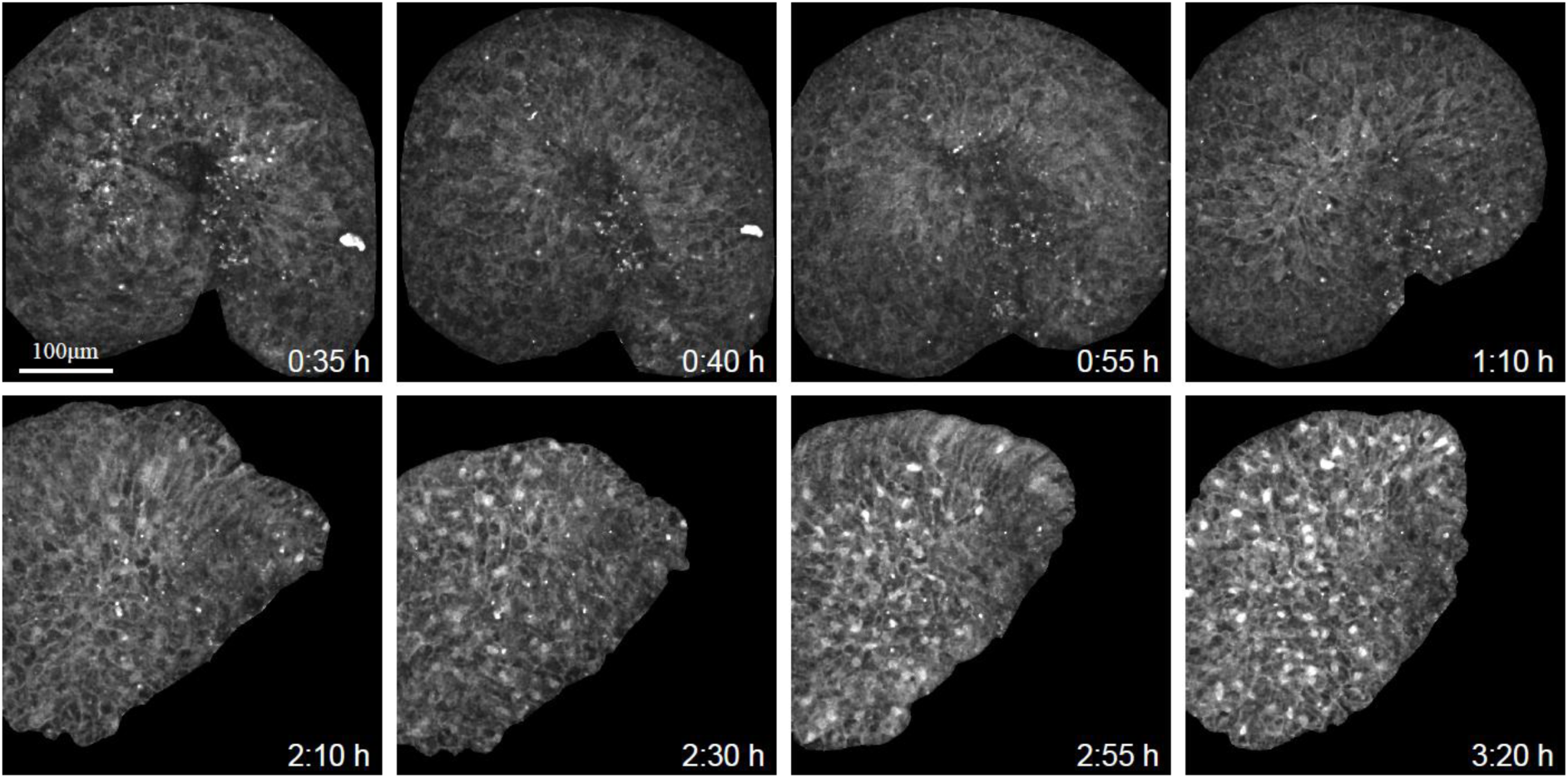
β-catenin dynamics during the initial folding of a regenerating tissue fragment. Spinning-disk confocal projected images from a time-lapse movie (Movie 5) of a regenerating tissue fragment in liquid medium. The upper row shows the initial folding and closure of the excised fragment into a spheroid. The lower row shows the subsequent emergence of the first widespread peak of nuclear β-catenin localization in the folded spheroid.

**Figure S4.**
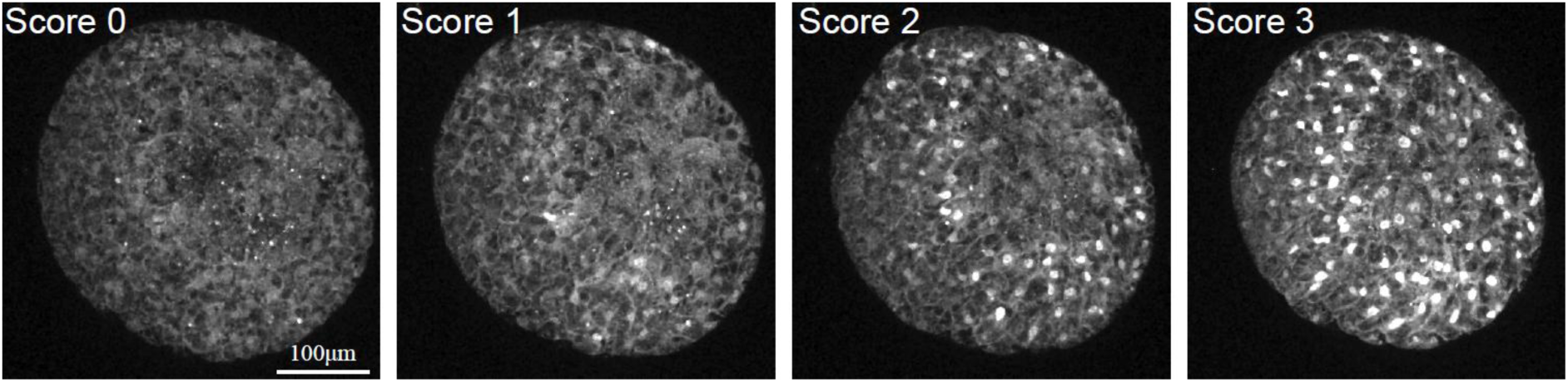
Manual scoring of nuclear β-catenin intensity. A regenerating spheroid expressing β-catenin-GFP in the ectoderm was imaged over time in an Airyscan microscope. Representative images from the time-lapse recording illustrate the four intensity scores: 0, no detectable nuclear fluorescence; 1, low nuclear fluorescence; 2, intermediate nuclear fluorescence; and 3, high nuclear fluorescence. The assigned score is indicated on each image. All images are displayed using identical brightness and contrast settings.

**Figure S5.**
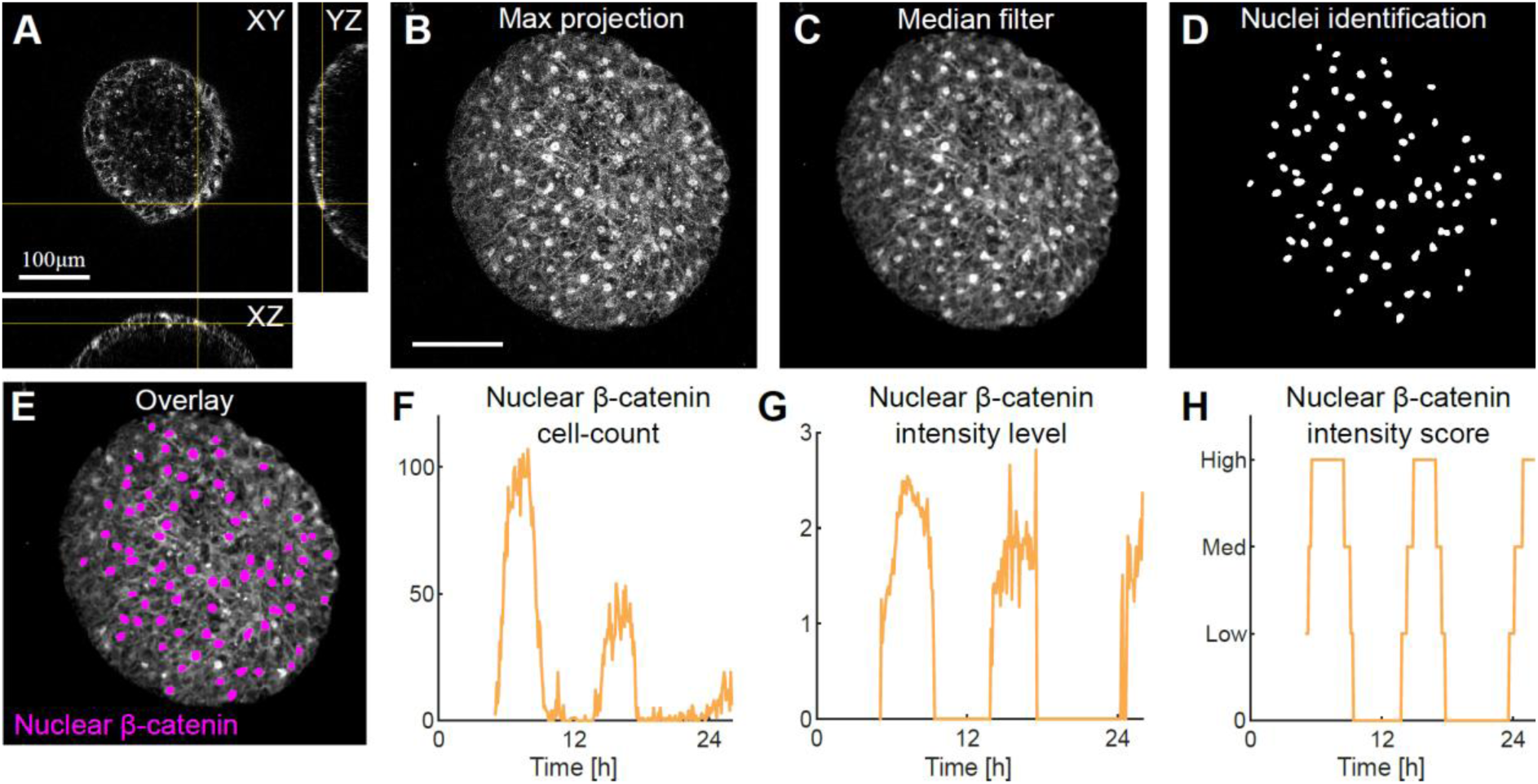
Automated analysis of β-catenin nuclear localization. A regenerating spheroid expressing β-catenin-GFP in the ectoderm was imaged over time in an Airyscan microscope. Nuclei with β-catenin localization are automatically identified (see Methods). (A) The images show cross-sections in the xy, xz, yz planes of the 3D image stack. (B,C) Raw (B) and median-filtered (C) maximum projection images. (D,E) Nuclei with β-catenin are automatically detected in the filtered projected image (Methods). The images show the mask of the detected nuclei (D) and a composite image showing the labeled nuclei (magenta) overlaid on the filtered projected image (E). (F,G) Graphs of the number (F) and mean intensity (G) of automatically detected nuclei with β-catenin localization over time in a regenerating tissue spheroid. The cell count is obtained by counting all identified nuclei and the intensity level is determined by averaging the mean signal intensity in the segmented nuclei. To reduce noise in the mean-intensity measurements when few nucleus-like objects are detected, measured intensity values are shown only for frames in which a three-frame moving average of the number of detected nuclei is at least 5. The plotted intensity is set to zero otherwise. (H) Graph showing the manually defined nuclear β-catenin intensity score over time in the same regenerating tissue spheroid (Fig. S4, Methods). The automated and manual analyses yield closely corresponding temporal profiles.

## Supplementary Movies

**Movie 1: β-catenin dynamics during head regeneration in a bisected *Hydra*.** Time-lapse, spinning-disk confocal movie of the oral end of a bisected *Hydra* expressing β-catenin-GFP in the ectoderm (Fig. 1D). Projected images of the β-catenin-GFP signal are shown. Nuclear β-catenin initially emerges across a broad region, subsequently declines, and later re-emerges within a smaller region at the regenerating head tip. The elapsed time from bisection is indicated, and the scale bar is 100 µm.

**Movie 2: β-catenin dynamics during foot regeneration in a bisected *Hydra*.** Time-lapse, spinning-disk confocal movie of the aboral end of a bisected Hydra expressing β-catenin-GFP in the ectoderm (Fig. 1E). Projected images of the β-catenin-GFP signal are shown. Nuclear β-catenin localization is not observed throughout foot regeneration. The elapsed time from bisection is indicated, and the scale bar is 100 µm.

**Movie 3: β-catenin dynamics in a regenerating *Hydra* tissue ring.** Time-lapse, spinning-disk confocal movie of an excised tissue ring expressing β-catenin-GFP in the ectoderm (Fig. 2E). Projected images of the entire ring are shown on the left, with rectangles indicating the oral (orange) and aboral (cyan) facing sides. Magnified views of these regions are shown on the right. Nuclei tracked manually are outlined in magenta in the magnified views. Nuclear β-catenin emerges on the oral-facing side, whereas no nuclear localization is detected on the aboral-facing side. The elapsed time from excision is indicated, and the scale bar is 100 µm.

**Movie 4: β-catenin dynamics in a regenerating *Hydra* tissue fragment viewed from the future head side.** Time-lapse, Airyscan confocal movie of a regenerating tissue fragment expressing β-catenin-GFP in the ectoderm (Fig. 3F). Projected images of the β-catenin-GFP signal are shown without annotation on the left and with an overlay indicating the automatically detected nuclei exhibiting β-catenin localization (magenta) on the right. Nuclear β-catenin initially increases across a broad region and then declines. A second, more spatially restricted increase is followed by another decline, after which nuclear β-catenin stabilizes within a focused region at the future head site. The elapsed time from excision is indicated, and the scale bar is 100 µm.

**Movie 5: β-catenin dynamics during the initial folding of a regenerating *Hydra* tissue fragment.** Time-lapse, spinning-disk confocal movie of an excised tissue fragment expressing β-catenin-GFP in the ectoderm and regenerating in liquid medium (Fig. S3). Projected images of the β-catenin-GFP signal show the fragment folding and closing into a spheroid, followed by the emergence of the first widespread peak of nuclear β-catenin localization. The elapsed time from excision is indicated, and the scale bar is 100 µm.

**Movie 6: β-catenin dynamics in a regenerating *Hydra* tissue fragment viewed from the future foot side.** Time-lapse, Airyscan confocal movie of a regenerating tissue fragment expressing β-catenin-GFP in the ectoderm (Fig. 3H). Projected images of the β-catenin-GFP signal are shown without annotation on the left and with an overlay indicating the automatically detected nuclei exhibiting β-catenin localization (magenta) on the right. Several nuclei with detectable β-catenin localization are present in the future foot region during the initial response. This nuclear signal subsequently disappears and does not re-emerge during the remainder of regeneration. The elapsed time from excision is indicated, and the scale bar is 100 µm.

## Notes

### Competing Interest Statement

The authors have declared no competing interest.

